# Comparative genomics analyses indicate differential methylated amine utilisation trait within members of the genus *Gemmobacter*

**DOI:** 10.1101/356014

**Authors:** E. Kröber, J Peixoto, MR Cunningham, L. Spurgin, D Wischer, R Kruger, D. Kumaresan

## Abstract

Methylated amines are ubiquitous in the environment and play a role in regulating the earth’s climate via a set of complex biological and chemical reactions. Microbial degradation of these compounds is thought to be a major sink. Recently we isolated a facultative methylotroph, *Gemmobacter* sp. LW-1, an isolate from the unique environment Movile Cave, Romania, which is capable of methylated amine utilisation as a carbon source. Here, using a comparative genomics approach, we investigate how widespread methylated amine utilisation trait is within the member of the bacterial genus *Gemmobacter*. Five genomes of different *Gemmobacter* species isolated from diverse environments, such as activated sludge, fresh water, sulphuric cave waters (Movile Cave) and the marine environment were available from the public repositories and used for the analysis. Our results indicate that some members of the genus *Gemmobacter*, namely *G. aquatilis, G. caeni* and *G*. sp. LW-1 have the genetic potential of methylated amine utilisation while others (*G. megaterium* and *G. nectariphilus*) have not.

## Introduction

Methylated amines (MAs) are ubiquitous in the environment with a variety of natural and anthropogenic sources including the oceans, vegetation, sediments and organic-rich soils, animal husbandry, food industry, pesticides, sewage, and automobiles, to mention only a few [1-3]. Methylated amines are also known to influence earth’s climate, via a series of complex biological and chemical interactions [4]. Some of the most abundant methylated amines found in the atmosphere are trimethylamine (TMA), dimethylamine (DMA) and monomethylamine (MMA) [1]. Microbial metabolism of methylated amines involves both aerobic and anaerobic microorganisms, e.g. some methanogenic archaea such as *Methanosarcina* and *Methanomicrobium* can use MAs to produce methane [5-7] while Gram-positive and Gram-negative methylotrophic bacteria can use MAs as carbon and nitrogen source [8]. Previously, MAs were typically associated with marine ecosystems as they are by-products of degradation of osmolytic chemicals such as glycine betaine, carnitine, choline and trimethylamine N-oxide [8]. However, recent studies have reported the detection and activity of aerobic methylotrophic bacteria that utilise MAs in a variety of natural and engineered environments [1, 9-12] and could play a major role in global C and N budgets.

Aerobic methylotrophs are a polyphyletic group of microorganisms capable of utilising one-carbon (C_1_) compounds such as methane, methanol or methylated amines as their sole source of carbon and energy [9, 13, 14]. Methylotrophs can degrade TMA to DMA by using the enzymes TMA dehydrogenase, TMA monooxygenase or TMA methyltransferase (under anaerobic conditions by methylotrophic methanogens), encoded by the genes *tdm, tmm* and *mtt*, respectively [15-18]. The enzymes DMA dehydrogenase (*dmd*) or DMA monooxygenase (*dmmDABC*) modulate the conversion of DMA to MMA [14, 15]. Two distinct pathways have been characterised for the oxidation of MMA [10]. The direct MMA-oxidation pathway mediated by a single enzyme (MMA dehydrogenase in Gram-negative bacteria and MMA oxidase in Gram-positive bacteria) converts MMA to formaldehyde and releases ammonium [19, 20]. The alternate pathway, referred to as the *N*-methylglutamate (NMG) pathway or indirect MMA-oxidation pathway, is mediated by three individual enzymes via the oxidation of MMA to gamma-glutamylmethylamide (GMA) and its further degradation to *N*-methylglutamate (NMG) and 5,10-methylenetetrahydrofolate (CH_2_ = H_4_F) [3, 10]. A stepwise conversion of MMA in the NMG pathway is modulated by the enzymes GMA synthetase (*gmaS*), NMG synthase (*mgsABC*) and NMG dehydrogenase (*mgdABCD*) [3, 8]. The capability to use MMA not only as a source for carbon but also for nitrogen is widespread in bacteria. Notably, the NMG pathway is not only restricted to methylotrophs but also present in non-methylotrophic bacteria that use MMA as a nitrogen but not as a carbon source [15, 21, 22].

In a recent study, we isolated an alphaproteobacterial facultative methylotrophic bacterium, *Gemmobacter* sp. LW-1 (recently renamed from *Catellibacterium* [23]) from the Movile Cave ecosystem (Mangalia, Romania) [24] that can use methylated amines as both carbon and nitrogen source [12] and subsequently obtained its genome sequence [25]. Using a ^13^C-MMA DNA based stable-isotope probing (SIP) experiment we also showed that *Gemmobacter* sp. LW-1 was indeed an active MMA utiliser in microbial mats from this environment [12]. This was the first report of methylated amine utilisation in a member of the bacterial genus *Gemmobacter*. However, growth on C_1_ compounds (methanol and formate) has been reported for the genus *Gemmobacter*, e.g. in *G. caeni* [26]. The genus *Gemmobacter* (family *Rhodobacteraceae*) currently comprises ten validated species: *Gemmobacter megaterium* [27], *G. nectariphilum* [23, 28], *G. aquatilis* [29], *G. caeni* [23, 26], *G. aquaticus* [23, 30], *G. nanjingense* [23, 31], *G. intermedius* [32], *G. lanyuensis* [33], *G. tilapiae* [34] and *G. fontiphilus* [23]. These species were isolated from a wide range of environments including fresh water environments (freshwater pond [29, 34], freshwater spring [23, 33]), coastal planktonic seaweed [27], white stork nestling [32], waste water and activated sludge [26, 28, 31], suggesting that members of the genus *Gemmobacter* are widely distributed in engineered and natural environments.

Here, using a comparative genomics approach we study how widespread methylated amine utilisation trait (i.e. metabolic potential) is within the members of the genus *Gemmobacter.* We used five isolate genomes for members within the genus *Gemmobacter* (*G*. sp. LW-1, *G. caeni, G. aquatilis, G. nectariphilus* and *G. megaterium)* alongside genomes of four closely related organisms within the family *Rhodobacteraceae* to show that the methylated amine utilisation trait is not common within the members of the genus *Gemmobacter.*

## Materials and Methods

### Genome data acquisition

Five *Gemmobacter* genomes (*G. caeni, G. aquatilis, G. nectariphilus, G. megaterium, Gemmobacter* sp. LW-1) available through the Integrated Microbial Genomes (IMG) database (https://img.jgi.doe.gov/) were used for comparative genome analysis [35]. Accession numbers and genome characteristics are listed in Supplementary Table S1.

### Phylogenetic analysis

Phylogenetic relatedness between the different members of the genus *Gemmobacter* was determined using phylogenetic trees constructed from 16S rRNA gene sequences (nucleotide) and metabolic gene sequences (*gmaS* and *mauA*; amino acids) involved in MMA utilisation. RNAmmer [36] was used to retrieve 16S rRNA gene sequences from the genome sequences. Multiple sequence alignment of 16S rRNA gene sequences was performed using the SINA alignment service via SILVA [37, 38] and subsequently imported into MEGA7 [39] to construct a maximum-likelihood nucleotide-based phylogenetic tree [40]. Bootstrap analysis was performed with 1000 replicates to provide confidence estimates for phylogenetic tree topologies [41].

To determine phylogenetic affiliations for the protein encoding genes *gmaS* and *mauA*, gene sequences retrieved from the genome sequences were aligned to homologous sequences retrieved from the NCBI Genbank database using Basic Local Alignment Search Tool (BLAST, blastx) [42] and curated *gmaS* sequences used for primer design in our previous study [12]. Amino acid sequences were aligned in MEGA7 [39] using ClustalW [43] and the alignment was subsequently used to construct maximum likelihood phylogenetic trees based on the JTT matrix-based model [44]. Bootstrap analysis was performed with 1000 replicates to provide confidence estimates for phylogenetic tree topologies [41].

We investigated the phylogeny of nine bacterial species – *Gemmobacter aquatilis, Gemmobacter caeni, Gemmobacter sp. LW-1, Gemmobacter megabacterium* DSM-26375, *Gemmobacter nectariphilus* DSM-15620, *Paracoccus denitrificans* PD1222, *Citreicella* sp. SE45, *Roseovarius* sp. TM1035 and *Rhodobacter sphaeroides* 241 – using the protein sequences of 831 orthologs present in single copy in all species identified using OrthoFinder [45] with default settings. For each set of orthologous proteins, multiple alignments were produced using MAFFT [46] with the --auto settings. Subsequently, conserved alignment blocks were identified using trimal [47] with the option -automated1. The phylogenetic reconstruction analysis was based in a final concatenated alignment consisting of 265,536 amino acid positions, which was the input for RAxML [48] with the option PROTCATLGF. The selection of best amino acid substitution model was performed after implementing the script ProteinModelSelection.pl provided by the RAxML package. In order to assess the environmental distribution of the genus *Gemmobacter*, we used MAPseq [49] (https://beta.microbeatlas.org) to survey the relative abundance based on 16S rRNA gene sequences (query sequence: *Gemmobacter aquatilis* DSM3857 (NR_104740.1 at 97% cut-off).

### Comparative genomic analyses

CGView Comparison Tool (CCT) was used to visually compare the genomes within the genus *Gemmobacter* [50]. CCT utilises BLAST to compare the genomes and the BLAST results are presented in a DNA-based graphical map [50]. Average Nucleotide Identity (ANI) [51] between different genomes was estimated using one-way ANI (best hit) and two-way ANI (reciprocal best hit) based on Goris *et al*. [52]. In addition the whole-genome based average nucleotide identity (gANI) and the p_r_^intra-species^ value were determined for *G.* sp. LW-1 and *G. caeni* (these two genomes revealed the closest ANI) based on Konstantinidis and Tiedje [53] via the Joint Genome Institute (JGI) platform (https://ani.jgi-psf.org/html/home.php; Version 0.3, April 2014). In order to determine if two genomes belong to the same species, the computation of empirical probabilities (p_r_^intra-species^) can be calculated as follows,

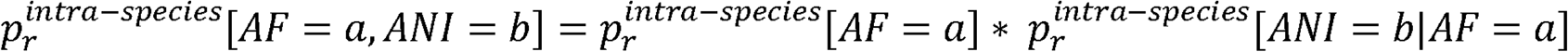

AF represents alignment fraction.

Pan-genome analysis including average amino acid identity (AAI) analysis, pan-genome tree construction and determination of core and dispensable genes and singletons (unique genes) was carried out using the Efficient Database framework for comparative Genome Analyses using BLAST score Ratios (EDGAR) platform [54]. Estimation of genomic completeness and contamination was carried out using the CheckM program [55].

In order to compare the genetic potential for methylated amine utilisation within the available *Gemmobacter* genomes, known protein sequences involved in methylated amine utilisation pathways [3, 15] were used as query sequences through the BLAST (blastp) program [42] available within the Rapid Annotation using Subsystem Technology (RAST) server [56]. The list of protein queries used is given in Supplementary Table S2.

## Results and discussion

### Analysis of phylogenetic relatedness

The phylogenetic relatedness of the five members within the genus *Gemmobacter* (*G.* sp. LW-1, *G. caeni, G. aquatilis, G. nectariphilus* and *G. megaterium*) was resolved based on 16S rRNA gene sequences (Figure 1). Three members of the genus *Gemmobacter* (*G.* sp. LW-1, *G. caeni*, and *G. aquatilis)* clustered together with several other related *Gemmobacter* and *Rhodobacter* 16S rRNA gene sequences retrieved from fresh water, soil and sediment and activated sludge environments (Figure 1). *G. nectariphilus* and *G. megaterium* sequences clustered together with *Paracoccus kawasakiensis* and other related *Gemmobacter* sequences from marine, fresh water and activated sludge environments (Figure 1). Based on the 16S rRNA gene sequences retrieved from public database and MAPseq analyses, we observed that the members of the genus *Gemmobacter* are widely distributed in engineered (such as activated sludge and clinical environments) and natural environments i.e. fresh water, soil and sediment, and marine environments (Figure 1 and Supplementary Figure S3). Phylogenomic analysis based on single copy marker genes (Supplementary Figure S4) revealed that *G.* sp LW1, *G. caeni and G. aquatilis* were closely related to *Rhodobacter sphaeroides* 241 whereas *G. megabacterium and G. nectariphilus* to *Paracoccus denitrificans.*

**Figure 1.**
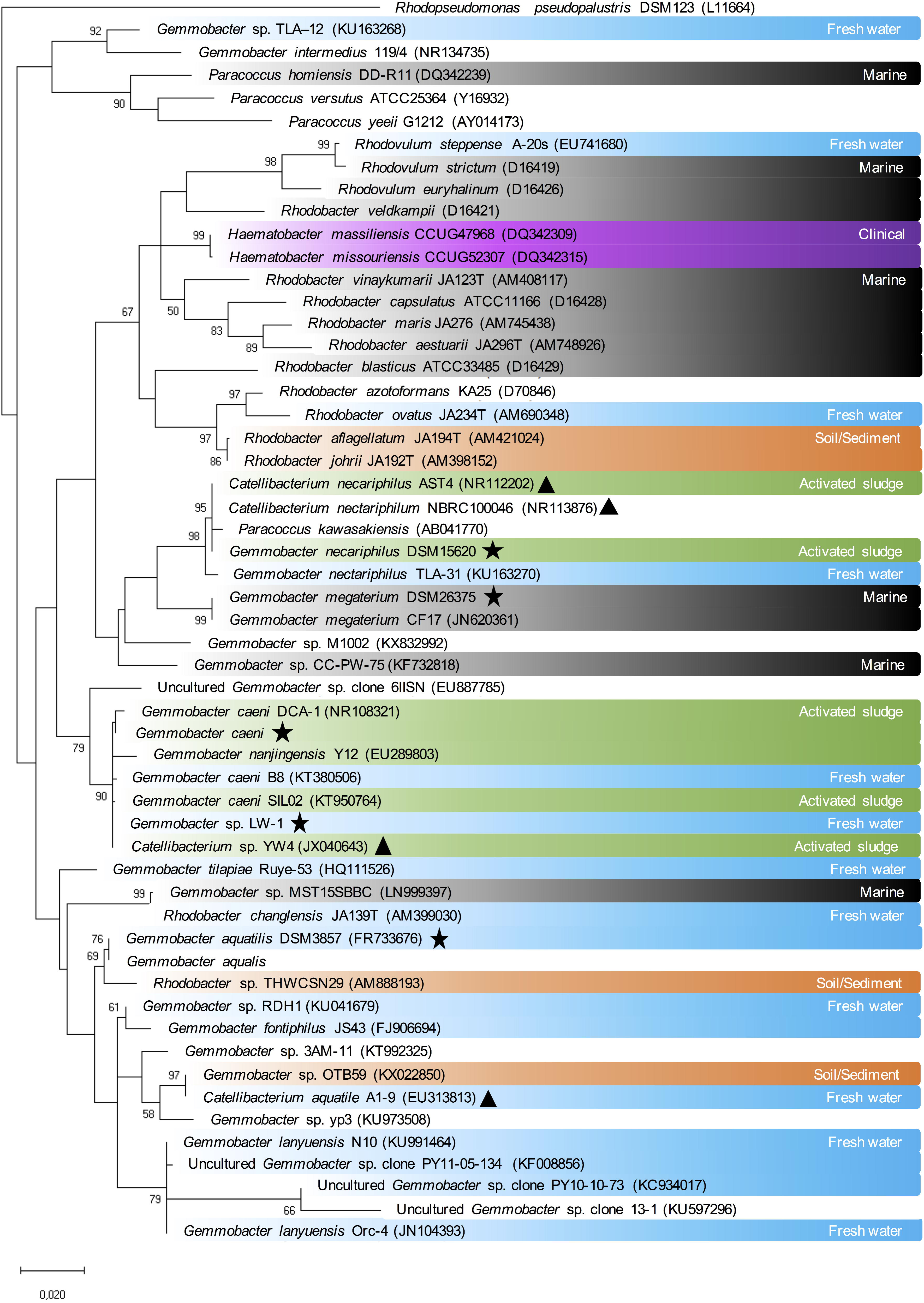
Phylogenetic tree based on 16S rRNA gene sequences. The tree was constructed using the maximum likelihood method for clustering and the Tamura-Nei model for computing evolutionary distances. Numbers at branches are bootstrap percentages >50% of 1000 replicates. Star represents the *Gemmobacter* species used for comparative genome analysis. Coloured boxes represent the habitat where the sequence was retrieved: blue (fresh water), orange (soil and sediment), green (activated sludge), grey (marine), purple (clinical source). Triangles represent sequences that are listed as *Catellibacterium* in the NCBI database, which have been recently reclassified to *Gemmobacter* [23]. Scale bar: 0.02 substitutions per nucleotide position.

GMA synthetase, a key enzyme in the NMG pathway, is encoded by the gene *gmaS. gmaS* sequences retrieved from the isolate genomes along with other ratified *gmaS* sequences were used to construct an amino acid-based phylogenetic tree (Figure 2). *gmaS* gene sequences retrieved from genomes of *G.* sp. LW-1, *G. caeni* and *G. aquatilis* clustered within Group I of alphaproteobacterial *gmaS* sequences containing sequences from marine and non-marine bacteria within the orders *Rhodobacterales* and *Rhizobiales* as described in Wischer et al. [12] and were closely related to *Paracoccus yeei, P.* sp. 1W-5 and *Rhodobacter* sp. 1W-5 (Figure 2). Whilst *gmaS* gene sequences were detected in three of the five investigated *Gemmobacter* genomes, *mauA* gene sequences were identified only in the genomes of *G. caeni* and *G.* sp. LW-1 (Supplementary Figure S1). It has been suggested that the NMG pathway for MMA utilisation is more universally distributed and more abundant across proteobacterial methylotrophs than the direct MMA oxidation pathway [57]. However, it should be noted that genes encoding for the enzymes within the NMG pathway (*gmaS*) can not only be detected in methylotrophs but also in non-methylotrophic bacteria that use MMA as a nitrogen source, but not as a carbon source [12, 15].

**Figure 2.**
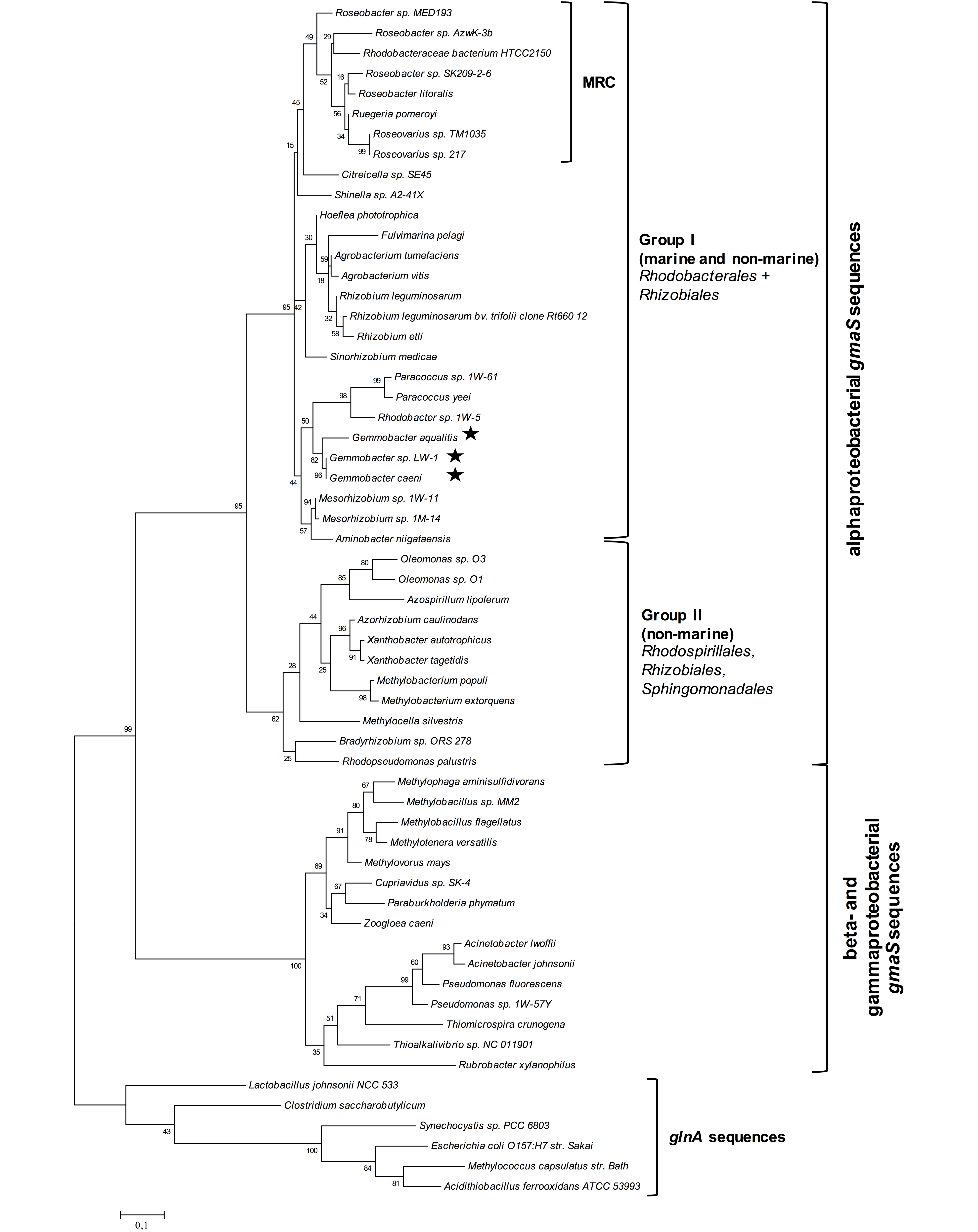
Maximum-likelihood phylogenetic tree based on *gmaS* sequences. The tree was constructed using amino acid sequences (GmaS) using the maximum-likelihood method based on the JTT matrix-based model. Members of the genus *Gemmobacter* used for genome comparison are represented with a star. Numbers at branches are bootstrap percentages >50% of 1000 replicates. Amino acid sequences of the glutamine synthetase type III (GlnA) were used as out-group. Scale bar: 0.1 substitutions per amino acid position. MRC, marine *Roseobacter* clade.

### A comparative genome analysis of members within the genus Gemmobacter

At the time of the analysis, five *Gemmobacter* genomes obtained from isolates from different environments were available (Figure 1 and Table 2). *Gemmobacter* genome sizes range from ∼3.96 Mb to ∼5.14 Mb with GC contents between 64.71% to 66.19% and genome completeness between 98.61% to 99.69% (Table 2; Supplementary Table 1). Analysis of sequence annotations revealed that on average 91.13% of the genomes consist of coding sequences.

The genomes were compared using the CGView comparison tool [50] (Figure 3). *Gemmobacter* sp. LW-1, isolated from the Movile Cave ecosystem was used as the reference genome and the results of the BLAST comparison with other *Gemmobacter* genomes are represented as a BLAST ring for each genome (Figure 3). Similarities between segments of the reference genome sequence and the other genome sequences are shown by a coloured arc beneath the region of similarity indicating the percentage of similarity as a colour code. Our analysis (Figure 3) revealed low identity levels (mostly <88%) between *Gemmobacter* sp. LW-1 and *G. aquatilis, G. nectariphilus* and *G. megaterium* across the genomes. Moreover, the analysis suggested several sites of potential insertion/deletion events in the genome of *Gemmobacter* sp. LW-1. Possible insertion/deletion regions can be identified as those gaps in the map where no homology is detected. For example, the region between 2200-2300 kbp (Figure 3) where a gap can be found in the otherwise contiguous homologous regions between the reference genome *G*. sp. LW-1 and the first of the query genomes (*G. caeni*). This might likely be due to a lack of hits or hits with low identity that can be spurious matches. Since it covers a large region we could possibly rule out that it is not an artefact arising from a lack of sensitivity in the BLAST analysis. Even though the genomes of *G.* sp. LW-1 and *G. caeni* are closely related, our analysis demonstrates that their genomes are not completely identical. Despite the fact that the majority of their genomes indicate very high identity levels (mostly >96-98% as shown by the dominance of dark red colours of the circle representing the BLAST hit identity between *G.* sp. LW-1 and *G. caeni*), many segments appear to be exclusive to *G.* sp. LW-1.

**Figure 3.**
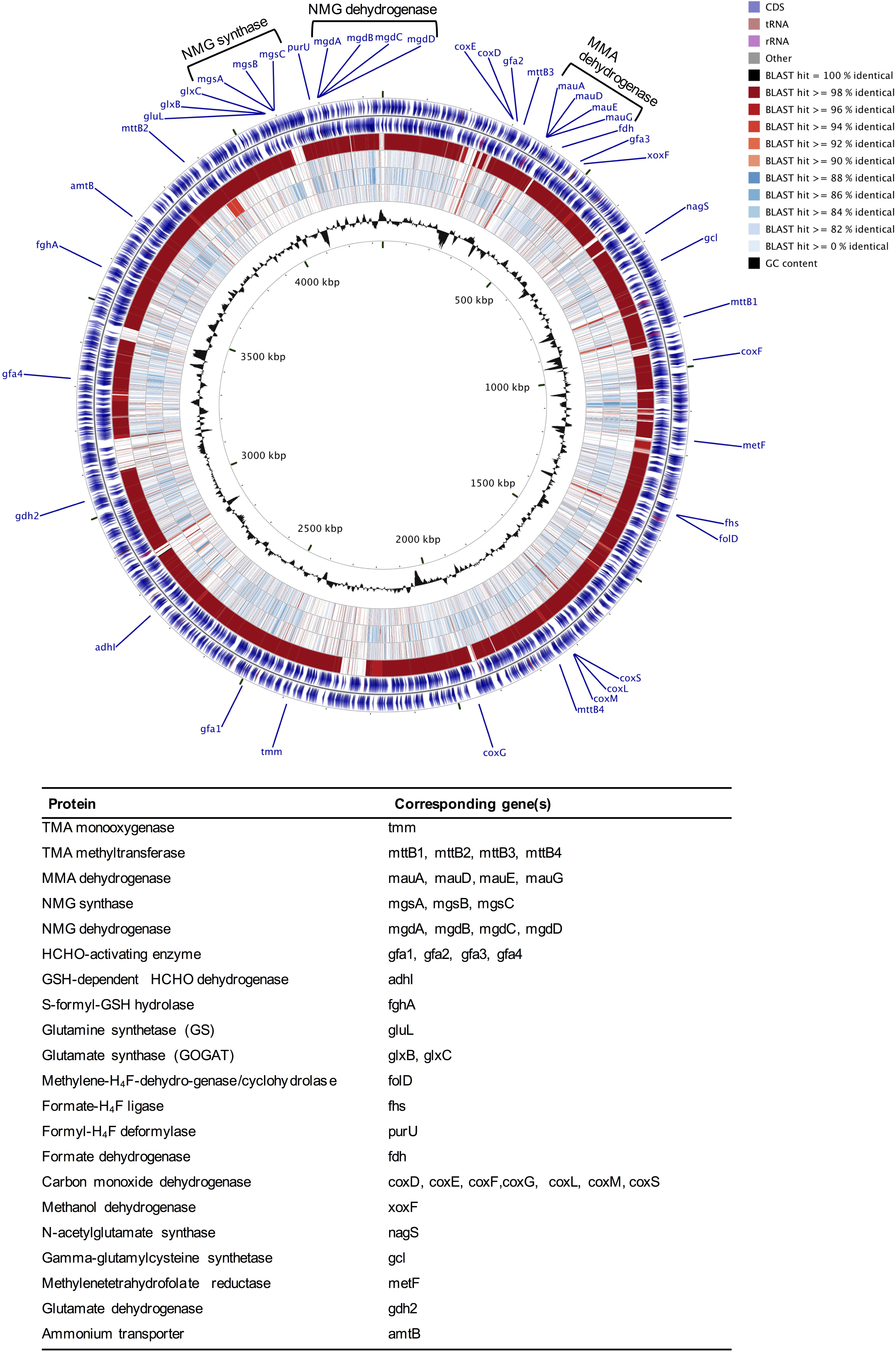
DNA BLAST map of *Gemmobacter* genomes. *Gemmobacter* sp. LW-1 was used as a reference genome against *Gemmobacter megaterium* (inner ring), *Gemmobacter aquatilis* (second inner ring), *Gemmobacter nectariphilus* (third ring), and *Gemmobacter caeni* (fourth ring). The fifth and sixth ring (outer rings) represent the CDS (blue), tRNA (maroon), and rRNA (purple) on the reverse and forward strand, respectively. The colour scale (inset) shows the level of sequence identity with the respective sequences from *G. megaterium, G. aquatilis, G. nectariphilus* and *G. caeni*. The locations of genes involved in methylotrophy are indicated at the outside of the map.

In order to further resolve the similarity between these genomes we calculated the average nucleotide identity (ANI) [51] (Supplementary Table S3 and Supplementary Figure S2A-D) and the average amino acid identity,(AAI; Supplementary Figure S2E). It is generally accepted that an ANI value of >95-96% can be used for species delineation [58, 59]. Our analysis revealed that *Gemmobacter* sp. LW-1 and *Gemmobacter caeni* share an ANI value of 98.62 (Supplementary Table S3) implying that both are in fact the same species. The genome-based average nucleotide identity (gANI) between *G.* sp. LW-1 and *G. caeni* was calculated as 98.70. The AF was calculated to be 0.91, which would result in a computed probability of 0.98 suggesting that both genomes might belong to the same species. However, it should be noted that these are draft genomes and a more in depth characterization of their physiology and phenotype is required to delineate these organisms at the level of strain.

Pan-genome analysis, carried out using the EDGAR platform [54], identified metabolic genes present in all *Gemmobacter* species (core genes), two or more *Gemmobacter* species (accessory or dispensable genes), and unique *Gemmobacter* species (singleton genes). A pan-genome tree was constructed (Figure 4A) based on the pan-genome dataset and neighbor-joining method [40]. As with the 16S-rRNA gene based phylogenetic tree (Figure 1), the five *Gemmobacter* species formed two main clusters in the pan-genome tree analysis (Figure 4A). The pan-genome tree also confirmed the phylogenetic closeness between *Gemmobacter caeni* and *Gemmobacter* sp. LW-1 (Figure 4A). According to pan-genome analysis of the five *Gemmobacter* genomes, a total of 9,286 genes were identified, consisting of 1,806 core genes, 3085 dispensable genes and 305, 1,072, 896, 1,165 and 957 singletons for *G*. sp. LW-1, *G. caeni, G. aquatilis, G. nectariphilus* and *G. megaterium*, respectively (Figure 4B).

**Figure 4.**
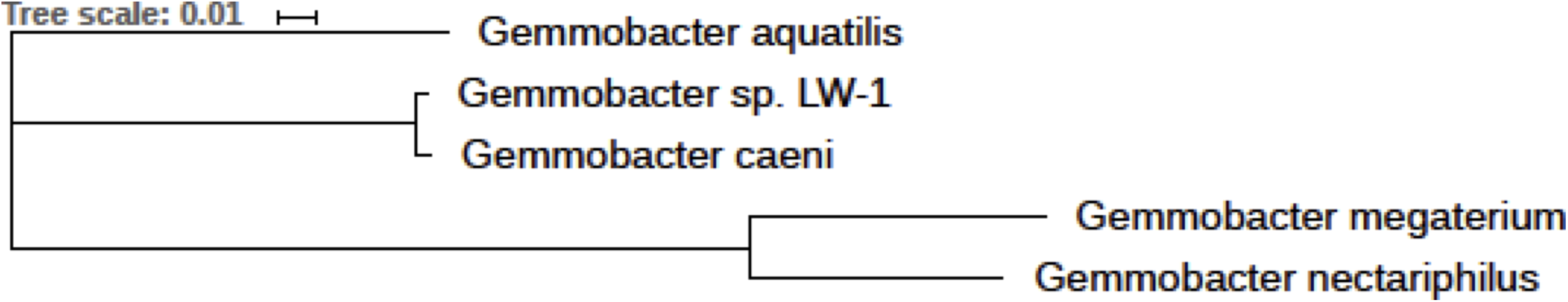
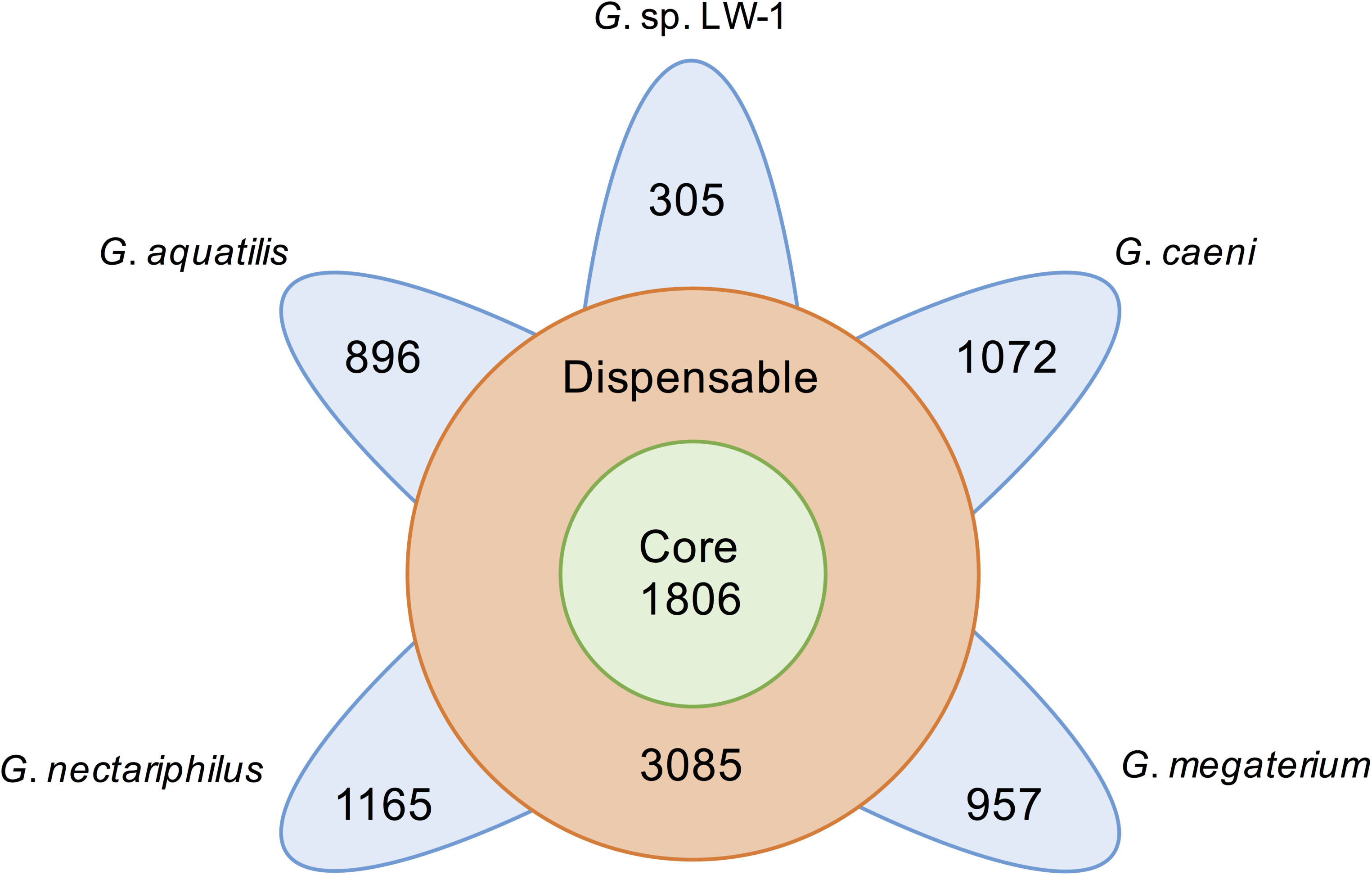
Pan-genome analysis. **(A)** Pan-genome tree consisting of five *Gemmobacter* species was constructed using the neighbour-joining method within the EDGAR platform. **(B)** Number of core, dispensable, and specific genes (singletons) of each *Gemmobacter* species.

### Methylated amine utilisation, N assimilation and C_1_ oxidation

Investigation of the methylated amine utilisation pathways in five *Gemmobacter* species revealed the presence of the genes encoding enzymes TMA dehydrogenase (*tmd*), TMA monooxygenase (*tmm*), TMAO demethylase (*tdm*) and DMA monooxygenase in genomes of *G*. sp. LW-1, *G. caeni* and *G. aquatilis* while none of these genes were detected in *G. nectariphilus* or *G. megaterium* (Figure 5). These findings are supported by results from a previous study which showed growth of *G.* sp. LW-1 on TMA as a carbon and nitrogen source [12]. *G*. sp. LW-1, *G. caeni* and *G. aquatilis* could potentially use the TMA oxidation pathway to convert TMA to DMA. Based on the genome sequences, it can be suggested that these three species could use the enzyme DMA monooxygenase (*dmmDABC*) to oxidize DMA to MMA but not the DMA dehydrogenase since the corresponding protein encoding gene (*dmd*) was not found (Figure 5).

**Figure 5.**
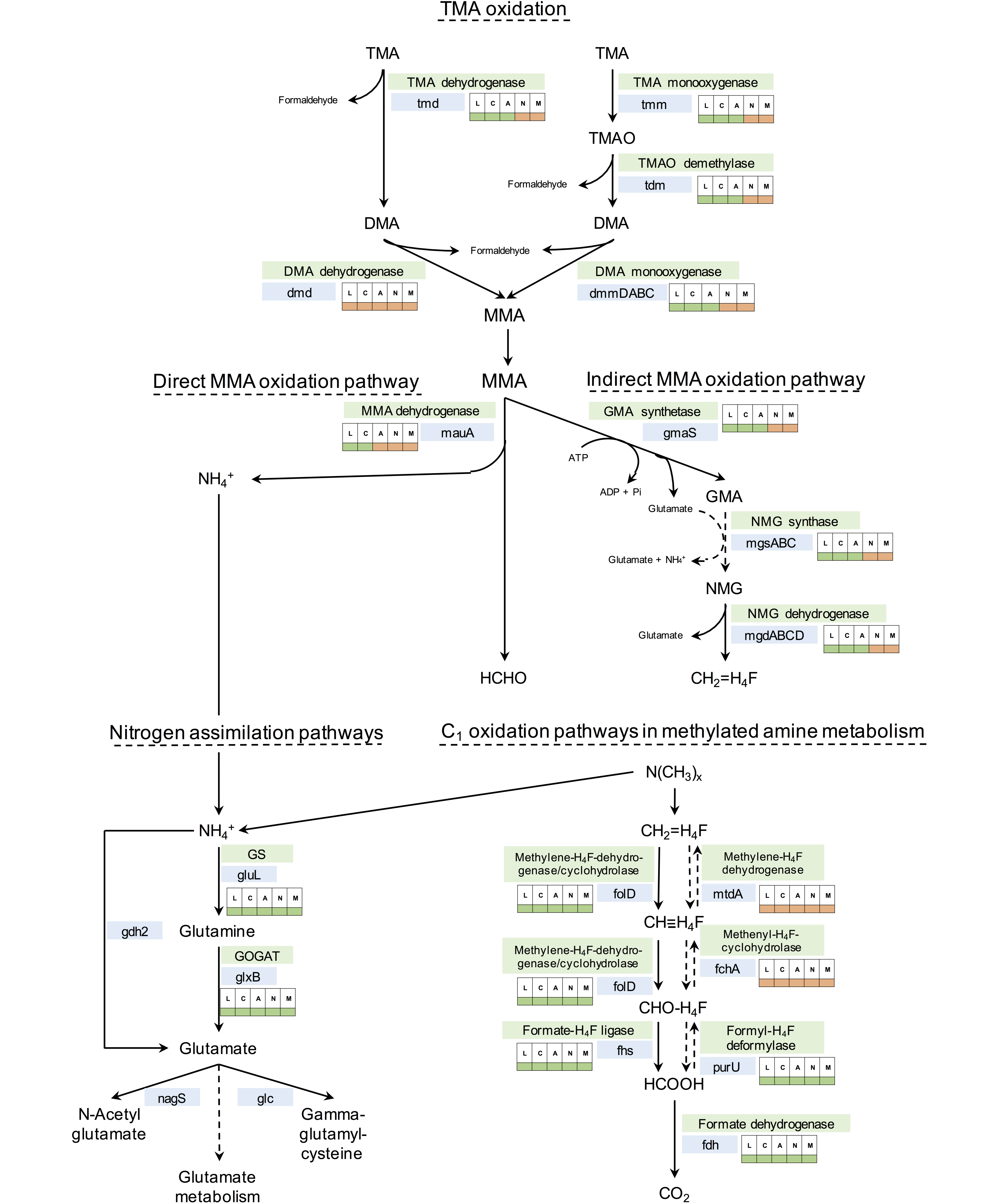
Metabolic pathways involved in methylated amine utilisation and one-carbon utilisation annotated with presence/absence of specific genes in the genomes of *Gemmobacter*. The analysis was based on a five-way comparison among *Gemmobacter* sp. LW-1 (L), *Gemmobacter caeni* (C), *Gemmobacter aquatilis* (A), *Gemmobacter nectariphilus* (N) and *Gemmobacter megaterium* (M). The color-coded boxes next to the genes indicate the presence (green) or absence (orange) of a gene in each genome.

We also compared the distribution of the direct MMA-oxidation and the NMG pathways in the genomes of five *Gemmobacter* species (Figure 5) and gene arrangement (Supplementary Figure S5). The direct MMA-oxidation pathway (*mauA*-dependent) is so far only known to be present in methylotrophic bacteria that can use MMA as a carbon source. Whereas the NMG pathway (*gmaS*-dependent) has been shown to be present in non-methylotrophic bacteria that can use MMA as a nitrogen source [12, 21, 57, 60]. Analysis of the genome sequences revealed that both *G*. sp. LW-1 and *G. caeni* possess genes for both MMA oxidation pathways (Figure 5). We have previously shown that *Gemmobacter* sp. LW-1 can use MMA and TMA as both a carbon and nitrogen source [12]. Genome sequence of *G. aquatilis* indicated the presence of genes involved only in the NMG pathway. In the facultative methylotroph *Methylobacterium extorquens* AM1 it has been shown that the NMG pathway is advantageous compared to the direct MMA-oxidation pathway [60]. NMG pathway enables facultative methylotrophic bacteria to switch between using MMA as a nitrogen source or as a carbon and energy source whereas the direct MMA oxidation pathway allows for rapid growth on MMA only as the primary energy and carbon source [60]. This could suggest that *G. aquatilis* might use the NMG pathway for utilising MMA as both nitrogen and carbon source. However, growth assays are required to confirm whether *G. aquatilis* can use MMA as a carbon source. We did not detect genes for either MMA oxidation pathways in the genome sequences of *G. nectariphilus* and *G. megaterium* suggesting the lack of genetic potential of these organisms to use MMA as either C or N source.

The C_1_ units derived from methylated amines need to be further oxidized when the nitrogen is sequestered without assimilation of the carbon from the methylated amines. Genome analysis confirmed that all five *Gemmobacter* species possess the genetic capability for C_1_ oxidation and also indicate that tetrahydrofolate (H_4_F) is the C_1_ carrier (Figure 5). The bifunctional enzyme 5,10-methylene-tetrahydrofolate dehydrogenase/ cyclohydrolase, encoded by the gene *folD*, was detected in all the *Gemmobacter* genomes (Figure 5/ Table 1). Genes encoding key enzymes in the C_1_ oxidation pathway via tetrahydromethanopterin (H_4_MPT) were not detected. [10]. The formate-tetrahydrofolate ligase, encoded by the gene *fhs* (Figure 5), provides C_1_ units for biosynthetic pathways [15]. However, the oxidation of formyl-H_4_F (CHO-H_4_F) can also be facilitated by *purU*, the gene encoding for the formyl-H_4_F deformylase. The formate dehydrogenase (*fdh*) mediates the last step of the C_1_ oxidation pathway, the oxidation of formate to CO_2_. The genes for the C_1_ oxidation pathway via H_4_F were detected in all five *Gemmobacter* genomes.

**Table 1.**
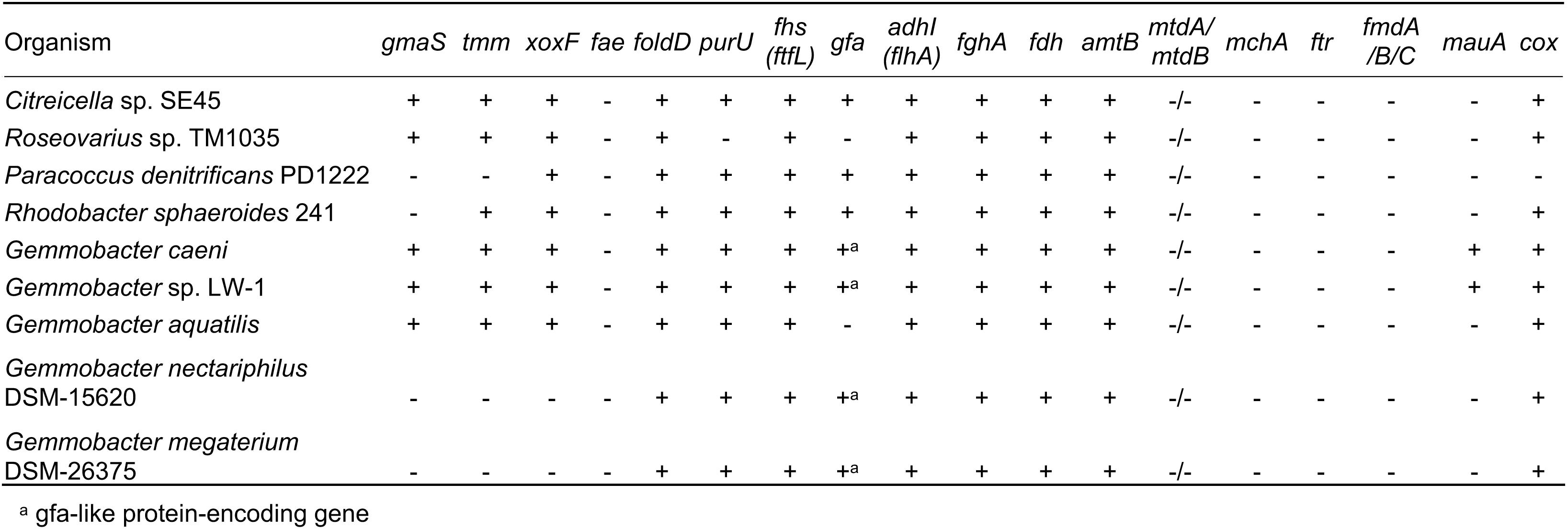
Comparative genomic analysis of methylated amine-utilising genes in genomes-sequenced *Gemmobacters* in comparison to selected marine *Roseobacter* clade bacteria. Shown is the presence (+) or absence (-) of specific genes in the genome sequences.

The *fae* gene, encoding the formaldehyde-activating enzyme that catalyses the reduction of formaldehyde with H_4_MPT was not detected in any of the five *Gemmobacter* genomes confirming that these members of the genus *Gemmobacter* lack the H_4_MPT pathway for formaldehyde oxidation (Figure 5/ Table 1). Investigation of the nitrogen assimilation pathway revealed the presence of the genes encoding glutamine synthetase (GS; *gluL*) and glutamine synthase (GOGAT; *glxB*) in all five *Gemmobacter* genomes. In bacteria this pathway is essential for glutamate synthesis at low ammonium concentrations [15].

Using comparative genome analysis we provide genome-based evidence that the two *Gemmobacter* isolates *G.* sp. LW-1 and *G. caeni* are capable of generating energy from complete oxidation of methylated amines via the H_4_F-dependent pathway using either the NMG pathway or the direct MMA oxidation pathway. *Gemmobacter aquatilis* is genetically capable of methylated amine degradation to yield formaldehyde and only encodes the genes for the NMG pathway, which indicates that *G. aquatilis* could use this pathway to use MMA as a nitrogen source. Both *G. nectariphilus* and *G. megaterium* genomes indicate the lack of potential to use methylated amines (Figure 5/Table 1).

*Gemmobacter* sp. LW-1 was isolated from the Movile Cave ecosystem [12]. Microbial mats and lake water within the cave have been shown to harbor a wide diversity of methylated amine-utilising bacteria [12, 61]. Whilst the mechanism of MAs production within the system has to be elucidated, it can be speculated that degradation of floating microbial mats (i.e. organic matter) could result in MAs [12]. Similarly, *G. caeni* isolated from activated sludge [26] could possibly use the MAs generated from organic matter degradation. Interestingly, whilst *G. megaterium* was isolated from a marine environment (seaweed [27]) possibly encountering MAs from the degradation of osmolytes such as glycine betaine (*N,N,N*-trimethylglycine) we did not detect metabolic genes involved in methylated amine utilisation. Our genome-based analyses suggest that the potential for methylated amine utilisation is differential within the members of the genus *Gemmobacter* that requires confirmation using growth/activity assays.

## Conclusions

In summary, three of the five investigated *Gemmobacter* genomes (*G.* sp. LW-1, *G. caeni* and *G. aquatilis*) indicated metabolic potential to utilise methylated amines, of which only two (*G.* sp. LW-1 and *G. caeni*) possess the genes for both MMA oxidation pathways, the NMG pathway and the direct MMA oxidation pathway. *G.* sp. LW-1 and *G. caeni* are facultative methylotrophs which could potentially use these pathways to utilise MMA as both a carbon and nitrogen source, while potentially *G. aquatilis* could only use the NMG pathway as a nitrogen source. Furthermore, the genomes of *G.* sp. LW-1 and *G. caeni* showed a high similarity to each other (>98%) suggesting that both belong to the same species. *G. megaterium* and *G. nectariphilus* genomes indicated no metabolic potential to utilise MAs. Phylogenetic, pan-genome and ANI analyses revealed that *G.* sp. LW-1 and *G. caeni* are closely related, although they were isolated from different environments. Overall, our results suggest that the trait for methylated amine utilisation could be independent from the habitat and localised factors or selection pressures could influence the ability of these organisms to use methylated amines. Access to *Gemmobacter* isolates with or without the genetic potential for methylated amine utilisation trait will allow us to perform physiological experiments in future to test how this trait can affect fitness of closely related organisms.

## Acknowledgements

We thank Dr Felipe Hernandes Coutinho (Federal University of Rio de Janeiro) for a fruitful discussion on genomic regions of insertion and deletion in *Gemmobacter* sp. LW-1. The EDGAR platform is financially supported by the BMBF grant FKZ 031A533 within the de.NBI network. This manuscript has been published as preprint at the preprint server for biology (bioRxiv.org, Kröber et al., 2018).

## Figures and Tables

**Supplementary Figure S1.** Maximum-likelihood phylogenetic tree (JTT matrix-based) of *mauA* sequences. Sequences from the genus *Gemmobacter* are marked with a star. Amino acid sequences (MauA) were aligned using the ClustalW algorithm. Numbers at branches are bootstrap percentages >50% of 1000 replicates. Scale bar: 0.1 substitutions per amino acid. Coloured boxes indicate *Alphaproteobacteria (*yellow), *Gammaproteobacteria (*red) and *Betaproteobacteria (*black).

**Supplementary Figure S2.** (A-D) Average nucleotide identity (ANI) analysis of *Gemmobacter* sp. LW-1 and *Gemmobacter caeni, Gemmobacter aquatilis, Gemmobacter nectariphilus* and *Gemmobacter megaterium* and (E) AAI analysis between those species

**Supplementary Figure S3.** Distribution of *Gemmobacter* in different ecosystems

**Supplementary Figure S4**. Maximum-likelihood phylogenetic tree of concatenated single copy marker genes from genomes of *Gemmobacter* and closely related organisms.

**Supplementary Figure S5**. Arrangement of genes involved in methylated amine utilisation

**Supplementary Table S1.** Genome characteristics of the five *Gemmobacter* isolate genomes used in this study.

**Supplementary Table S2.** List of protein queries used for the genome comparison with their accession number.

**Supplementary Table S3.** Average nucleotide identity (ANI) values between *Gemmobacter* sp. LW-1 and *Gemmobacter caeni, Gemmobacter aquatilis, Gemmobacter nectariphilus, Gemmobacter megaterium, Rhodobacter sphaeroides* and *Paracoccus denitrificans*

## REFERENCES

1. Ge X, Wexler AS, Clegg SL (2011) Atmospheric amines – Part I. A review. Atmospheric Environment 45: 524–546. doi:http://dx.doi.org/10.1016/j.atmosenv.2010.10.012

2. Schade GW, Crutzen PJ (1995) Emission of Aliphatic-Amines from Animal Husbandry and Their Reactions - Potential Source of N2o and Hcn. J Atmos Chem 22: 319–346. doi: Doi 10.1007/Bf00696641

3. Latypova E, Yang S, Wang YS, Wang T, Chavkin TA, Hackett M, Schäfer H, Kalyuzhnaya MG (2010) Genetics of the glutamate-mediated methylamine utilization pathway in the facultative methylotrophic beta-proteobacterium Methyloversatilis universalis FAM5. Molecular microbiology 75: 426–439.

4. Carpenter LJ, Archer SD, Beale R (2012) Ocean-atmosphere trace gas exchange. Chemical Society Reviews 41: 6473–6506. doi: 10.1039/c2cs35121h

5. Lyimo TJ, Pol A, Jetten MSM, Op den Camp HJM (2009) Diversity of methanogenic archaea in a mangrove sediment and isolation of a new Methanococcoides strain. Fems Microbiol Lett 291: 247–253. doi: 10.1111/j.1574-6968.2008.01464.x

6. Burke SA, Lo SL, Krzycki JA (1998) Clustered genes encoding the methyltransferases of methanogenesis from monomethylamine. J Bacteriol 180: 3432–3440.

7. Liu Y, Whitman WB (2008) Metabolic, phylogenetic, and ecological diversity of the methanogenic archaea. Ann N Y Acad Sci 1125: 171–189. doi: 10.1196/annals.1419.019

8. Chen Y, Scanlan J, Song L, Crombie A, Rahman MT, Schäfer H, Murrell JC (2010) γ-Glutamylmethylamide is an essential intermediate in the metabolism of methylamine by Methylocella silvestris. Applied and environmental microbiology 76: 4530–4537.

9. Chistoserdova L, Kalyuzhnaya MG, Lidstrom ME (2009) The Expanding World of Methylotrophic Metabolism. Annual Review of Microbiology 63: 477–499. doi: 10.1146/annurev.micro.091208.073600

10. Chistoserdova L (2011) Modularity of methylotrophy, revisited. Environmental Microbiology 13: 2603–2622.

11. Chen Y, Wu L, Boden R, Hillebrand A, Kumaresan D, Moussard H, Baciu M, Lu Y, Colin Murrell J (2009) Life without light: microbial diversity and evidence of sulfur- and ammonium-based chemolithotrophy in Movile Cave. ISME J 3: 1093–1104.

12. Wischer D, Kumaresan D, Johnston A, El Khawand M, Stephenson J, Hillebrand-Voiculescu AM, Chen Y, Colin Murrell J (2015) Bacterial metabolism of methylated amines and identification of novel methylotrophs in Movile Cave. ISME J 9: 195–206. doi: 10.1038/ismej.2014.102

13. Anthony C (1982) The biochemistry of methylotrophs. Academic Press, London

14. Lidstrom ME (2006) Aerobic Methylotrophic Prokaryotes. In: Dworkin, M, Falkow, S, Rosenberg, E, Schleifer, K-H, Stackebrandt, E (eds.) The Prokaryotes: Volume 2: Ecophysiology and Biochemistry. Springer New York, New York, NY, pp. 618–634

15. Chen Y (2012) Comparative genomics of methylated amine utilization by marine Roseobacter clade bacteria and development of functional gene markers (tmm, gmaS). Environmental Microbiology 14: 2308–2322.

16. Lidbury I, Murrell JC, Chen Y (2014) Trimethylamine N-oxide metabolism by abundant marine heterotrophic bacteria. Proceedings of the National Academy of Sciences 111: 2710–2715.

17. Paul L, Ferguson DJ, Krzycki JA (2000) The trimethylamine methyltransferase gene and multiple dimethylamine methyltransferase genes of Methanosarcina barkeri contain in-frame and read-through amber codons. J Bacteriol 182: 2520–2529.

18. Lidbury I, Mausz MA, Scanlan DJ, Chen Y (2017) Identification of dimethylamine monooxygenase in marine bacteria reveals a metabolic bottleneck in the methylated amine degradation pathway. ISME J 11: 1592–1601. doi: 10.1038/ismej.2017.31

19. McIntire WS, Wemmer DE, Chistoserdova A, Lidstrom ME (1991) A new cofactor in a prokaryotic enzyme: tryptophan tryptophylquinone as the redox prosthetic group in methylamine dehydrogenase. Science 252: 817–824.

20. Chistoserdova AY, Christoserdova LV, McIntire WS, Lidstrom ME (1994) Genetic organization of the mau gene cluster in Methylobacterium extorquens AM1: complete nucleotide sequence and generation and characteristics of mau mutants. J Bacteriol 176: 4052–4065.

21. Chen Y, McAleer KL, Murrell JC (2010) Monomethylamine as a nitrogen source for a nonmethylotrophic bacterium, Agrobacterium tumefaciens. Applied and environmental microbiology 76: 4102–4104.

22. Taubert M, Grob C, Howat AM, Burns OJ, Pratscher J, Jehmlich N, von Bergen M, Richnow HH, Chen Y, Murrell JC (2017) Methylamine as a nitrogen source for microorganisms from a coastal marine environment. Environmental Microbiology 19: 2246–2257. doi: 10.1111/1462-2920.13709

23. Chen WM, Cho NT, Huang WC, Young CC, Sheu SY (2013) Description of Gemmobacter fontiphilus sp nov., isolated from a freshwater spring, reclassification of Catellibacterium nectariphilum as Gemmobacter nectariphilus comb. nov., Catellibacterium changlense as Gemmobacter changlensis comb. nov., Catellibacterium aquatile as Gemmobacter aquaticus nom. nov., Catellibacterium caeni as Gemmobacter caeni comb. nov., Catellibacterium nanjingense as Gemmobacter nanjingensis comb. nov., and emended description of the genus Gemmobacter and of Gemmobacter aquatilis. International Journal of Systematic and Evolutionary Microbiology 63: 470–478. doi: 10.1099/ijs.0.042051-0

24. Kumaresan D, Wischer D, Stephenson J, Hillebrand-Voiculescu A, Murrell JC (2014) Microbiology of Movile Cave - A chemolithoautotrophic ecosystem. Geomicrobiology Journal 31: 186–193.

25. Kumaresan D, Wischer D, Hillebrand-Voiculescu AM, Murrell JC (2015) Draft Genome Sequences of Facultative Methylotrophs, Gemmobacter sp. Strain LW1 and Mesorhizobium sp. Strain 1M-11, Isolated from Movile Cave, Romania. Genome Announc 3. doi: 10.1128/genomeA.01266-15

26. Zheng J-W, Chen Y-G, Zhang J, Ni Y-Y, Li W-J, He J, Li S-P (2011) Description of Catellibacterium caeni sp. now., reclassification of Rhodobacter changlensis Anil Kumar et al. 2007 as Catellibacterium changlense comb. nov. and emended description of the genus Catellibacterium. International Journal of Systematic and Evolutionary Microbiology 61: 1921–1926.

27. Liu J-J, Zhang X-Q, Chi F-T, Pan J, Sun C, Wu M (2014) Gemmobacter megaterium sp. nov., isolated from coastal planktonic seaweeds. International Journal of Systematic and Evolutionary Microbiology 64: 66–71. doi: doi:10.1099/ijs.0.050955-0

28. Tanaka Y, Hanada S, Manome A, Tsuchida T, Kurane R, Nakamura K, Kamagata Y (2004) Catellibacterium nectariphilum gen. nov., sp. nov., which requires a diffusible compound from a strain related to the genus Sphingomonas for vigorous growth. International Journal of Systematic and Evolutionary Microbiology 54: 955–959. doi: doi:10.1099/ijs.0.02750-0

29. Rothe B, Fischer A, Hirsch P, Sittig M, Stackebrandt E (1987) The phylogenetic position of the budding bacteria Blastobacter aggregatus and Gemmobacter aquatilis gen., nov. sp. nov. Archives of Microbiology 147: 92–99. doi: 10.1007/bf00492911

30. Liu Y, Xu C-J, Jiang J-T, Liu Y-H, Song X-F, Li H, Liu Z-P (2010) Catellibacterium aquatile sp. nov., isolated from fresh water, and emended description of the genus Catellibacterium Tanaka et al. 2004. International Journal of Systematic and Evolutionary Microbiology 60: 2027–2031. doi: doi:10.1099/ijs.0.017632-0

31. Zhang J, Chen S-A, Zheng J-W, Cai S, Hang B-J, He J, Li S-P (2012) Catellibacterium nanjingense sp. nov., a propanil-degrading bacterium isolated from activated sludge, and emended description of the genus Catellibacterium. International Journal of Systematic and Evolutionary Microbiology 62: 495–499. doi: doi:10.1099/ijs.0.029819-0

32. Kämpfer P, Jerzak L, Wilharm G, Golke J, Busse H-J, Glaeser SP (2015) Gemmobacter intermedius sp. nov., isolated from a white stork (Ciconia ciconia). International Journal of Systematic and Evolutionary Microbiology 65: 778–783. doi: doi:10.1099/ijs.0.000012

33. Sheu S-Y, Shiau Y-W, Wei Y-T, Chen W-M (2013) Gemmobacter lanyuensis sp. nov., isolated from a freshwater spring. International Journal of Systematic and Evolutionary Microbiology 63: 4039–4045. doi: doi:10.1099/ijs.0.052399-0

34. Sheu S-Y, Sheu D-S, Sheu F-S, Chen W-M (2013) Gemmobacter tilapiae sp. nov., a poly-β-hydroxybutyrate-accumulating bacterium isolated from a freshwater pond. International Journal of Systematic and Evolutionary Microbiology 63: 1550–1556. doi: doi:10.1099/ijs.0.044735-0

35. Markowitz VM, Chen I-MA, Palaniappan K, Chu K, Szeto E, Pillay M, Ratner A, Huang J, Woyke T, Huntemann M (2013) IMG 4 version of the integrated microbial genomes comparative analysis system. Nucleic acids research 42: D560–D567.

36. Lagesen K, Hallin P, Rødland EA, Stærfeldt H-H, Rognes T, Ussery DW (2007) RNAmmer: consistent and rapid annotation of ribosomal RNA genes. Nucleic acids research 35: 3100–3108.

37. Pruesse E, Peplies J, Glöckner FO (2012) SINA: accurate high-throughput multiple sequence alignment of ribosomal RNA genes. Bioinformatics 28: 1823–1829.

38. Pruesse E, Quast C, Knittel K, Fuchs BM, Ludwig W, Peplies J, Glöckner FO (2007) SILVA: a comprehensive online resource for quality checked and aligned ribosomal RNA sequence data compatible with ARB. Nucleic acids research 35: 7188–7196.

39. Kumar S, Stecher G, Tamura K (2016) MEGA7: Molecular Evolutionary Genetics Analysis version 7.0 for bigger datasets. Molecular biology and evolution 33: 1870–1874.

40. Saitou N, Nei M (1987) The neighbor-joining method: a new method for reconstructing phylogenetic trees. Molecular biology and evolution 4: 406–425.

41. Felsenstein J (1985) Confidence limits on phylogenies: an approach using the bootstrap. Evolution 39: 783–791.

42. Altschul SF, Gish W, Miller W, Myers EW, Lipman DJ (1990) Basic local alignment search tool. Journal of molecular biology 215: 403–410.

43. Thompson JD, Higgins DG, Gibson TJ (1994) CLUSTAL W: improving the sensitivity of progressive multiple sequence alignment through sequence weighting, position-specific gap penalties and weight matrix choice. Nucleic acids research 22: 4673–4680.

44. Jones DT, Taylor WR, Thornton JM (1992) The rapid generation of mutation data matrices from protein sequences. Bioinformatics 8: 275–282.

45. Emms DM, Kelly S (2015) OrthoFinder: solving fundamental biases in whole genome comparisons dramatically improves orthogroup inference accuracy. Genome biology 16: 157. doi: 10.1186/s13059-015-0721-2

46. Nakamura T, Yamada KD, Tomii K, Katoh K (2018) Parallelization of MAFFT for large-scale multiple sequence alignments. Bioinformatics 34: 2490–2492. doi: 10.1093/bioinformatics/bty121

47. Capella-Gutiérrez S, Silla-Martínez JM, Gabaldón T (2009) trimAl: a tool for automated alignment trimming in large-scale phylogenetic analyses. Bioinformatics 25: 1972–1973.

48. Stamatakis A (2014) RAxML version 8: a tool for phylogenetic analysis and post–analysis of large phylogenies. Bioinformatics 30: 1312–1313.

49. Matias Rodrigues JF, Schmidt TSB, Tackmann J, von Mering C (2017) MAPseq: highly efficient k-mer search with confidence estimates, for rRNA sequence analysis. Bioinformatics 33: 3808–3810. doi: 10.1093/bioinformatics/btx517

50. L. Grant JR, Arantes AS, Stothard P (2012) Comparing thousands of circular genomes using the CGView Comparison Tool. BMC genomics 13: 202.

51. Rodriguez-R LM, Konstantinidis KT (2016) The enveomics collection: a toolbox for specialized analyses of microbial genomes and metagenomes. PeerJ Preprints.

52. Goris J, Konstantinidis KT, Klappenbach JA, Coenye T, Vandamme P, Tiedje JM (2007) DNA–DNA hybridization values and their relationship to whole-genome sequence similarities. International journal of systematic and evolutionary microbiology 57: 81–91.

53. Konstantinidis KT, Tiedje JM (2005) Genomic insights that advance the species definition for prokaryotes. P Natl Acad Sci USA 102: 2567–2572. doi: 10.1073/pnas.0409727102

54. Blom J, Kreis J, Spanig S, Juhre T, Bertelli C, Ernst C, Goesmann A (2016) EDGAR 2.0: an enhanced software platform for comparative gene content analyses. Nucleic Acids Research 44: W22–W28. doi: 10.1093/nar/gkw255

55. Parks DH, Imelfort M, Skennerton CT, Hugenholtz P, Tyson GW (2015) CheckM: assessing the quality of microbial genomes recovered from isolates, single cells, and metagenomes. Genome research: gr. 186072.186114.

56. Aziz RK, Bartels D, Best AA, DeJongh M, Disz T, Edwards RA, Formsma K, Gerdes S, Glass EM, Kubal M (2008) The RAST Server: rapid annotations using subsystems technology. BMC genomics 9: 75.

57. Nayak DD, Marx CJ (2015) Experimental horizontal gene transfer of methylamine dehydrogenase mimics prevalent exchange in nature and overcomes the methylamine growth constraints posed by the sub-optimal N-Methylglutamate pathway. Microorganisms 3: 60–79.

58. Richter M, Rossello-Mora R (2009) Shifting the genomic gold standard for the prokaryotic species definition. Proceedings of the National Academy of Sciences of the United States of America 106: 19126–19131.

59. Kim M, Oh H-S, Park S-C, Chun J (2014) Towards a taxonomic coherence between average nucleotide identity and 16S gene sequence similarity for species demarcation of prokaryotes. International Journal of Systematic and Evolutionary Microbiology 64: 346–351.

60. Nayak DD, Agashe D, Lee M-C, Marx CJ (2016) Selection maintains apparentlydegenerate metabolic pathwaysdueto tradeoffsin using methylamine for carbon versus nitrogen. Current Biology 26: 1–11.

61. Kumaresan D, Stephenson J, Doxey AC, Bandukwala H, Brooks E, Hillebrand-Voiculescu A, Whiteley AS, Murrell JC (2018) Aerobic proteobacterial methylotrophs in Movile Cave: genomic and metagenomic analyses. Microbiome 6. doi: ARTN 1 10.1186/s40168-017-0383-2

