## Supplementary figures and images for "Comparative genomics analyses indicate differential methylated amine utilisation trait within members of the genus *Gemmobacter*"

### Supplementary Figure 1

#### Supplementary Figure S1

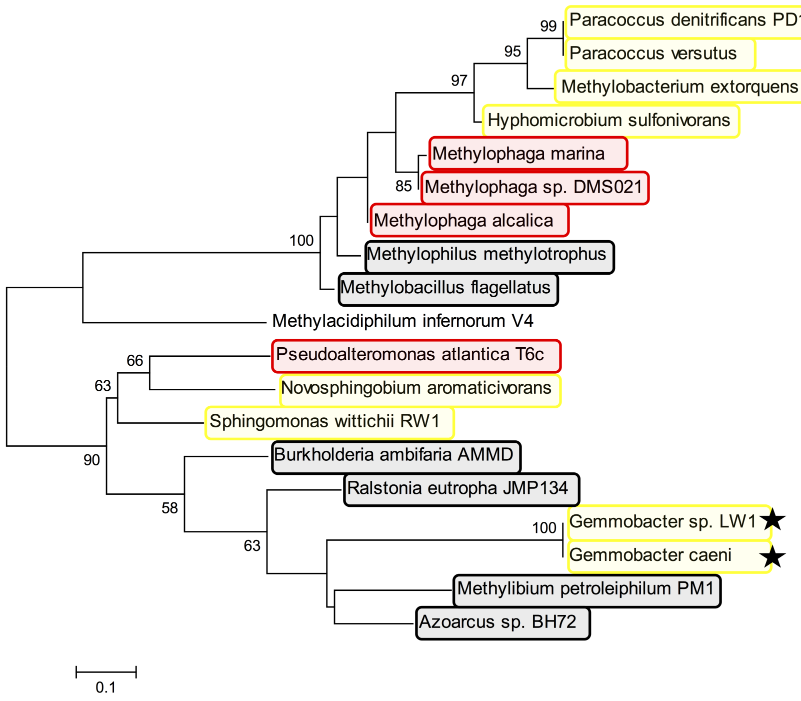

### Supplementary Figure 2

Supplementary Figure S2

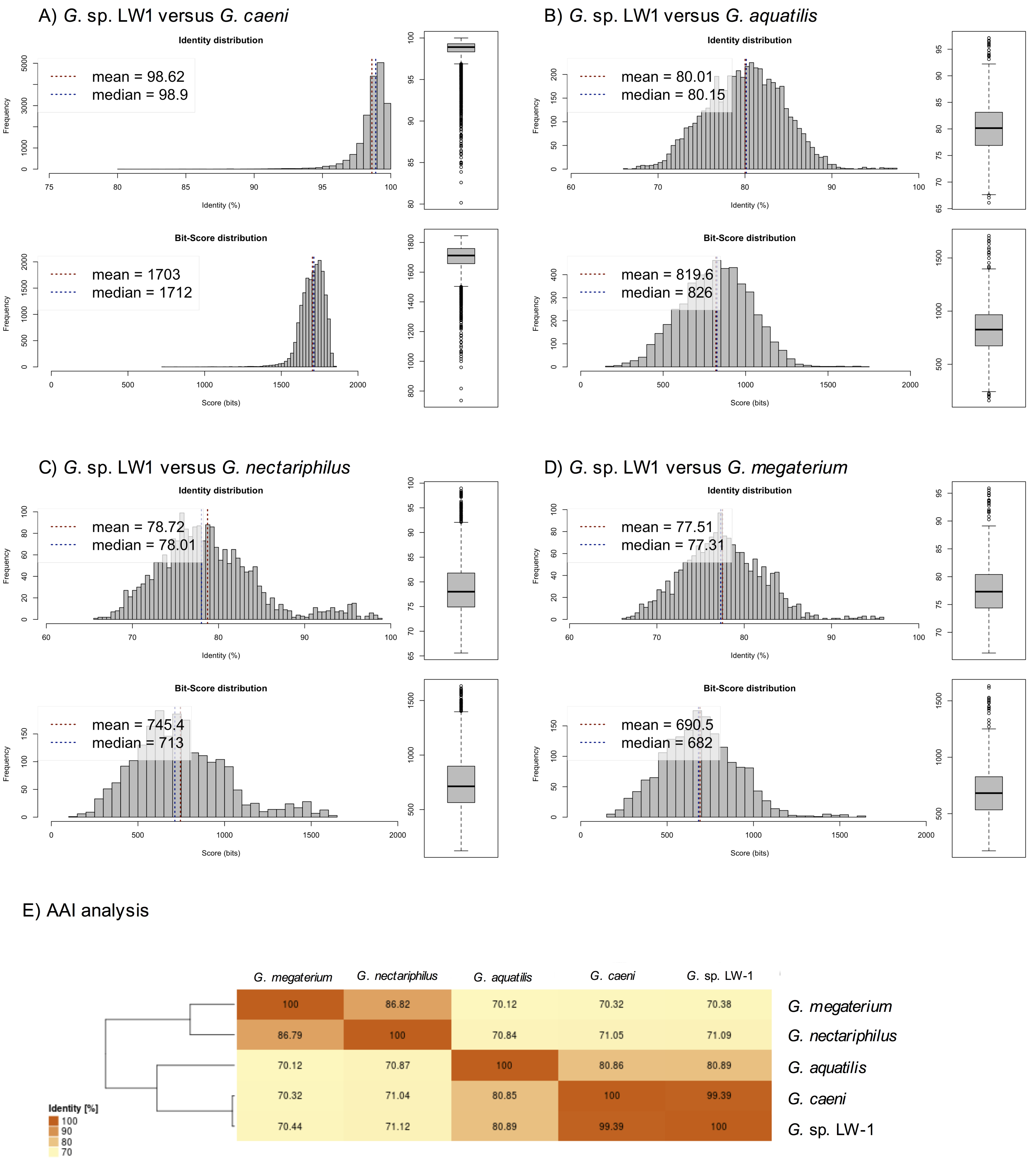

### Supplementary Figure 4

**Supplementary Figure S4**

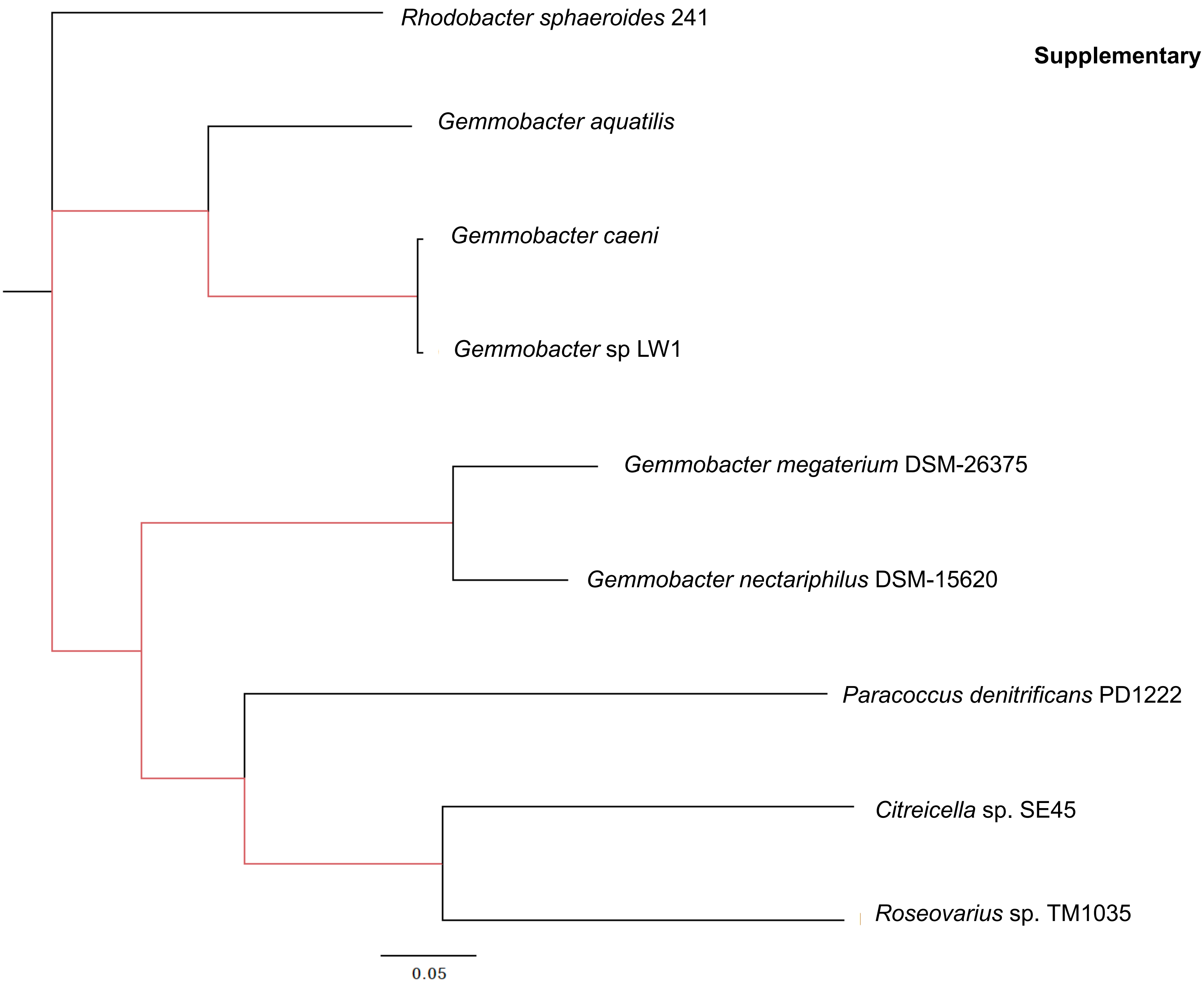

### Supplementary Figure 5

**Supplementary Figure S5**

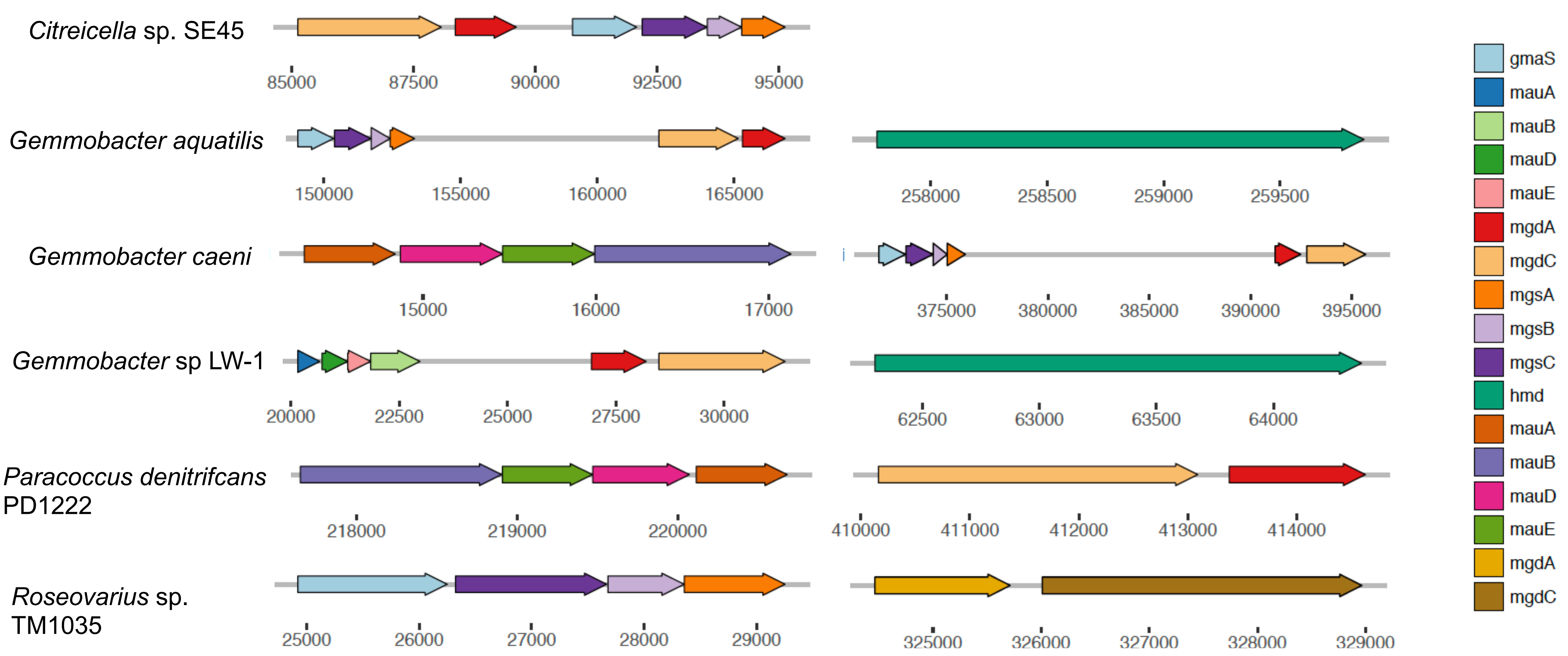
