## Supplementary Figure 3 for "Comparative genomics analyses indicate differential methylated amine utilisation trait within members of the genus *Gemmobacter*"

**Supplementary Figure S3**

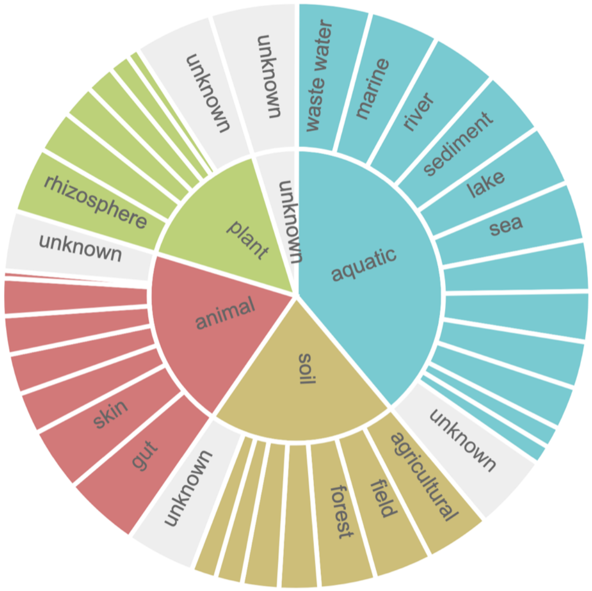


**aquatic: 4,810 samples**

- waste water: 7.71% (1,050)
- marine: 4.78% (650)
- river: 3.58% (486)
- sediment: 3.55% (483)
- lake: 2.27% (308)
- sea: 1.82% (248)
- groundwater: 0.82% (112)
- estuary: 0.80% (109)
- ocean: 0.57% (77)
- reservoir: 0.39% (53)
- brine: 0.05% (7)
- ice: 0.04% (6)
- unknown: 9.01% (1,220)

**soil: 1,540 samples**

- agricultural: 2.99% (406)
- field: 1.61% (219)
- forest: 1.38% (188)
- farm: 0.30% (41)
- paddy: 0.23% (31)
- desert: 0.09% (12)
- peatland: 0.07% (10)
- shrub: 0.01% (1)
- tundra: 0.01% (1)
- unknown: 4.64% (631)

**animal: 1,870 samples**

- gut: 7.96% (1,080)
- skin: 2.73% (371)
- gastric: 0.35% (48)
- urogenital: 0.30% (41)
- oral: 0.26% (35)
- lung: 0.24% (33)
- bone: 0.01% (2)
- unknown: 1.93% (262)

**plant: 2,040 samples**

- rhizosphere: 3.40% (462)
- leaf: 0.38% (51)
- seed: 0.16% (22)
- flower: 0.10% (14)
- wood: 0.05% (7)
- stem: 0.02% (3)
- sprout: 0.01% (1)
- unknown: 10.90% (1,480)

**unknown: 3,320 samples**

- unknown: 24.50% (3,320)
