## Supplementary Tables for "Comparative genomics analyses indicate differential methylated amine utilisation trait within members of the genus *Gemmobacter*"

### Slide 1
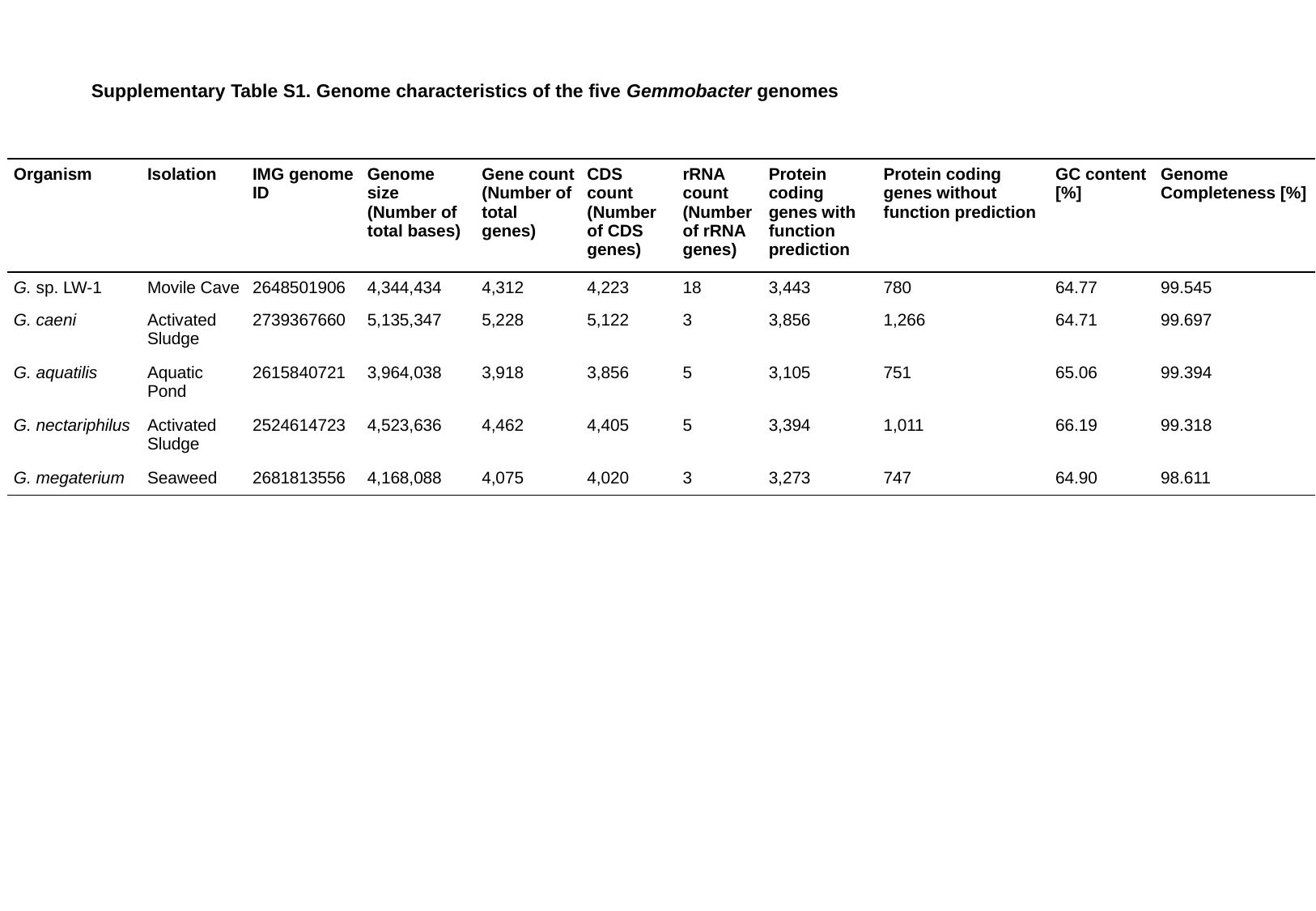

Supplementary Table S1. Genome characteristics of the five Gemmobacter genomes
| Organism | Isolation | IMG genome ID | Genome size (Number of total bases) | Gene count (Number of total genes) | CDS count (Number of CDS genes) | rRNA count (Number of rRNA genes) | Protein coding genes with function prediction | Protein coding genes without function prediction | GC content [%] | Genome Completeness [%] |
| --- | --- | --- | --- | --- | --- | --- | --- | --- | --- | --- |
| G. sp. LW-1 | Movile Cave | 2648501906 | 4,344,434 | 4,312 | 4,223 | 18 | 3,443 | 780 | 64.77 | 99.545 |
| G. caeni | Activated Sludge | 2739367660 | 5,135,347 | 5,228 | 5,122 | 3 | 3,856 | 1,266 | 64.71 | 99.697 |
| G. aquatilis | Aquatic Pond | 2615840721 | 3,964,038 | 3,918 | 3,856 | 5 | 3,105 | 751 | 65.06 | 99.394 |
| G. nectariphilus | Activated Sludge | 2524614723 | 4,523,636 | 4,462 | 4,405 | 5 | 3,394 | 1,011 | 66.19 | 99.318 |
| G. megaterium | Seaweed | 2681813556 | 4,168,088 | 4,075 | 4,020 | 3 | 3,273 | 747 | 64.90 | 98.611 |

### Slide 2
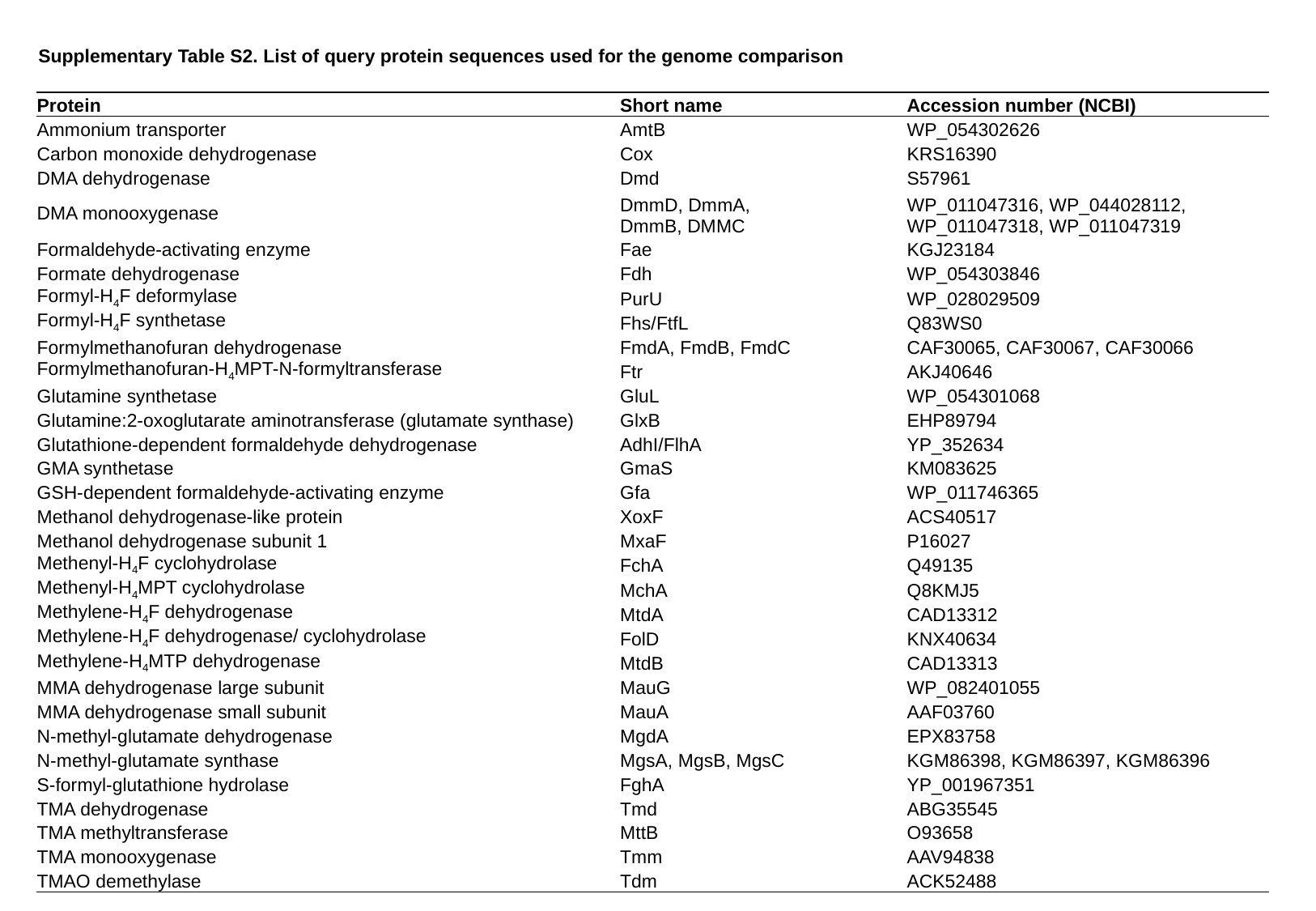

Supplementary Table S2. List of query protein sequences used for the genome comparison
| Protein | Short name | Accession number (NCBI) |
| --- | --- | --- |
| Ammonium transporter | AmtB | WP\_054302626 |
| Carbon monoxide dehydrogenase | Cox | KRS16390 |
| DMA dehydrogenase | Dmd | S57961 |
| DMA monooxygenase | DmmD, DmmA, DmmB, DMMC | WP\_011047316, WP\_044028112, WP\_011047318, WP\_011047319 |
| Formaldehyde-activating enzyme | Fae | KGJ23184 |
| Formate dehydrogenase | Fdh | WP\_054303846 |
| Formyl-H_4_F deformylase | PurU | WP\_028029509 |
| Formyl-H_4_F synthetase | Fhs/FtfL | Q83WS0 |
| Formylmethanofuran dehydrogenase | FmdA, FmdB, FmdC | CAF30065, CAF30067, CAF30066 |
| Formylmethanofuran-H_4_MPT-N-formyltransferase | Ftr | AKJ40646 |
| Glutamine synthetase | GluL | WP\_054301068 |
| Glutamine:2-oxoglutarate aminotransferase (glutamate synthase) | GlxB | EHP89794 |
| Glutathione-dependent formaldehyde dehydrogenase | AdhI/FlhA | YP\_352634 |
| GMA synthetase | GmaS | KM083625 |
| GSH-dependent formaldehyde-activating enzyme | Gfa | WP\_011746365 |
| Methanol dehydrogenase-like protein | XoxF | ACS40517 |
| Methanol dehydrogenase subunit 1 | MxaF | P16027 |
| Methenyl-H_4_F cyclohydrolase | FchA | Q49135 |
| Methenyl-H_4_MPT cyclohydrolase | MchA | Q8KMJ5 |
| Methylene-H_4_F dehydrogenase | MtdA | CAD13312 |
| Methylene-H_4_F dehydrogenase/ cyclohydrolase | FolD | KNX40634 |
| Methylene-H_4_MTP dehydrogenase | MtdB | CAD13313 |
| MMA dehydrogenase large subunit | MauG | WP\_082401055 |
| MMA dehydrogenase small subunit | MauA | AAF03760 |
| N-methyl-glutamate dehydrogenase | MgdA | EPX83758 |
| N-methyl-glutamate synthase | MgsA, MgsB, MgsC | KGM86398, KGM86397, KGM86396 |
| S-formyl-glutathione hydrolase | FghA | YP\_001967351 |
| TMA dehydrogenase | Tmd | ABG35545 |
| TMA methyltransferase | MttB | O93658 |
| TMA monooxygenase | Tmm | AAV94838 |
| TMAO demethylase | Tdm | ACK52488 |

### Slide 3
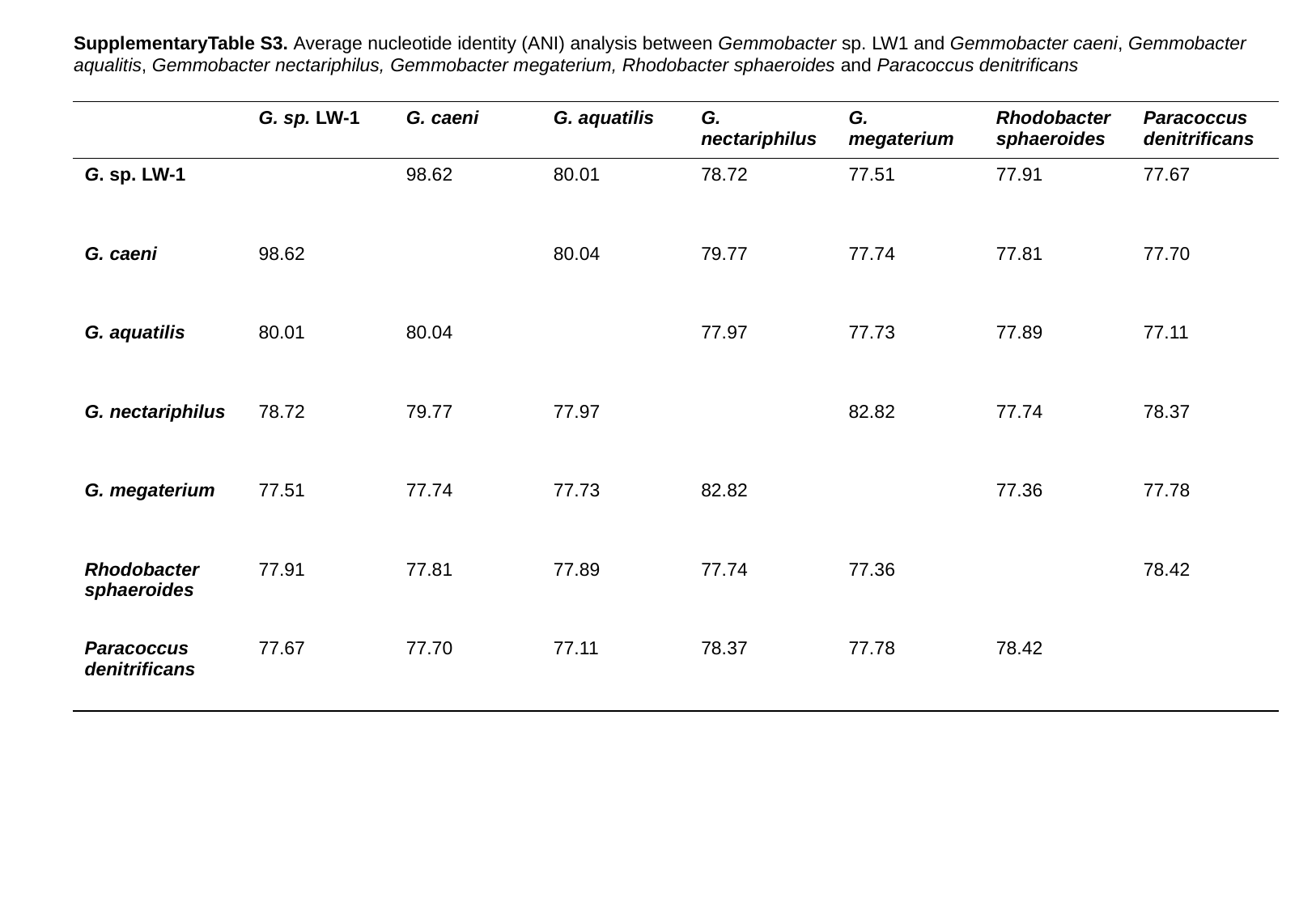

SupplementaryTable S3. Average nucleotide identity (ANI) analysis between Gemmobacter sp. LW1 and Gemmobacter caeni, Gemmobacter aqualitis, Gemmobacter nectariphilus, Gemmobacter megaterium, Rhodobacter sphaeroides and Paracoccus denitrificans
| | G. sp. LW-1 | G. caeni | G. aquatilis | G. nectariphilus | G. megaterium | Rhodobacter sphaeroides | Paracoccus denitrificans |
| --- | --- | --- | --- | --- | --- | --- | --- |
| G. sp. LW-1 | | 98.62 | 80.01 | 78.72 | 77.51 | 77.91 | 77.67 |
| G. caeni | 98.62 | | 80.04 | 79.77 | 77.74 | 77.81 | 77.70 |
| G. aquatilis | 80.01 | 80.04 | | 77.97 | 77.73 | 77.89 | 77.11 |
| G. nectariphilus | 78.72 | 79.77 | 77.97 | | 82.82 | 77.74 | 78.37 |
| G. megaterium | 77.51 | 77.74 | 77.73 | 82.82 | | 77.36 | 77.78 |
| Rhodobacter sphaeroides | 77.91 | 77.81 | 77.89 | 77.74 | 77.36 | | 78.42 |
| Paracoccus denitrificans | 77.67 | 77.70 | 77.11 | 78.37 | 77.78 | 78.42 | |
